# Engagement in science content via online video leveraging television science fiction (*Star Trek: Discovery*)

**DOI:** 10.1101/2023.03.12.532266

**Authors:** Mohamed A. F. Noor, Kim Manturuk

**Affiliations:** Duke University, USA

## Abstract

Using science fiction stories to teach real-world science concepts may improve student engagement and motivate creativity. Here, we explore the effectiveness of a science outreach video leveraging interest in science fiction television content at engaging an audience self-selected through online social media. We find that responding audience members report knowledge gains, that the science-fiction connection increased their interest, and that many may not have viewed this science content without the science-fiction connection. Overall, these findings support the use of fictional narrative connections to engage with non-student learners around science topics, extending the findings from classroom settings to online videos.

## INTRODUCTION

People appreciate narrative stories, which generally cover a temporal sequence of events that impact particular characters (e.g., Dahlstrom, 2014). Narrative story formats are commonly associated with fiction, both in print and in other modalities (Schank and Berman, 2002), but various scholars have explored the use of narratives as a means of engaging or persuading audiences about relevant natural or social scientific topics. Narrative formats have gained further recent support for science communication (e.g., National Academies of Sciences, 2017; Martinez-Conde and Macknik, 2017) particularly around topics controversial among the public such as global climate change and common ancestry of species. While narratives have been used in science communication primarily via historical or non-fiction stories (e.g., ElShafie, 2018; Solomon, 2002), some authors have argued that fictional narratives may also be effective if used ethically and if containing a basis in non-fictional elements (e.g., Dahlstrom and Ho, 2012; Dahlstrom, 2014; Avraamidou and Osborne, 2009) given consumers often retain and apply what they learn from such stories (e.g., Appel and Richter, 2007; Marsh et al., 2003).

Science fiction stories provide promising narratives because they are often grounded in real science. STEM professionals are even sometimes recruited to serve as consultants to collaborate with the fiction writers and producers to edit the content for accuracy (Loverd et al., 2018). The narratives of such science fiction stories help with student engagement, motivate creativity, and provide “a sense of meaning” (Vrasidas et al., 2015; Jordan and Silva, 2019). Critical analysis of such science fiction has numerous advantages (e.g., Gough, 1993; Barnett and Kafka, 2007), potentially even when there are errors that can be used as illustrative lessons regarding common misconceptions (Marsh et al., 2012).

However, most studies that we found assessing the impact of lessons incorporating science fiction on engagement and understanding focused on classroom settings. The public also consumes science content after the end of their formal schooling, sometimes through scholarly works aimed at them but arguably more often through representations in mass media. Unlike a classroom setting, in which students are a “captive audience” potentially through college requirements, the public have more freedom to choose which items to engage. This freedom provides further rationale for STEM professionals leveraging interest in popular culture elements such as science fiction as a venue for providing science education content to the broader public (e.g., Noor, 2018).

In this study, we explore the effectiveness of a science outreach video leveraging science fiction television content at engaging an audience self-selected through online social media. The main questions we explore are 1) whether respondents report that they learned from the video, 2) whether the science-fiction connection helped with their engagement & interest, and 3) whether the science fiction connection is broadening the viewership of this educational content. We also explore dependencies on how interesting they found the content relative to their familiarity with the franchise or specific shows, or with the timing or venue of their last formal science education. Importantly, we are not assessing whether this approach is “more” effective than a traditional science video—such an analysis would have been biased by the self-selection of the audience—but instead whether this approach is reaching and teaching at least some audience that would not have otherwise consumed the science content.

## MATERIALS AND METHODS

### Subjects

The data for this qualitative research were collected through an anonymous Qualtrics survey following informed consent. The link to the survey was posted to a variety of Star Trek-related online groups and discussion forums as well as the personal social media accounts of the researchers (see Supplemental Table 1). The survey link was also “shared” or “retweeted” by many others on social media. All posts directed people to a website where they were invited to watch a video and answer a few questions subsequently. The data collection protocol was approved by the Duke University IRB (protocol 2022-0265). The survey remained open from January 26, 2022, to April 18, 2022. During that time, 254 responses were recorded.

### Measures

Eight questions were asked with a 5-option Likert-like scale:

1. How much did you enjoy the video? (Not at all / a little / a moderate amount / a lot / a great deal)
2. How much do you feel you knew about prions prior to watching this video? (Not at all / a little / a moderate amount / a lot / a great deal)
3. How much do you feel you know about prions now? (Not at all / a little / a moderate amount / a lot / a great deal)
4. Did the Star Trek connection make this video more engaging to you? (A lot less / a little less / no difference / a little more / a lot more)
5. Did watching this video increase your knowledge about prions? (Not at all / a little / a moderate amount / a lot / a great deal)
6. How likely would you be to watch a science video about prions that did not relate to Star Trek? (Extremely unlikely / somewhat unlikely / neither likely nor unlikely / somewhat likely / extremely likely)
7. How did you feel about the length of this video for conveying the science covered? (Much too short / somewhat too short / right length / somewhat too long / much too long)
8. How much of the science do you feel you understood from what was presented in this video? (Not at all / a little / a moderate amount / a lot / a great deal)

Five additional questions were also asked:

9. How often do you watch science videos or read about science (outside the context of science fiction)? (Never / a few times a year / once or twice a month / around once a week / several times a week or more often)
10. Where did you last take a formal science class? (Never / elementary or middle school / high school / 2-year college or training program / military / 4-year college / university graduate program / online platform)
11. How long ago was that class? (Taking now / within past year / 1-5 years ago / 5-10 years ago / more than 10 years ago)
12. Were you familiar with the Star Trek Discovery episode that this video discussed? (Not familiar with Star Trek / familiar with Star Trek but not Discovery / familiar with Discovery but not this episode / familiar with this episode but not prion connection / familiar with the episode and remember prions being part of story)
13. Before you watched the video for this study, had you watched it elsewhere such as on YouTube? (No / think watched before but didn’t remember details / watched it and remembered it)

### Procedures

After informed consent, subjects were instructed to watch a 7-minute video (https://youtu.be/kBGwj9ZInis) explaining “prions” and contextualizing with their usage in an episode of *Star Trek: Discovery* (season 3, episode 5, “Die Trying”). Subjects included in the analysis then submitted responses to the 13 questions described above. Because the video had been available online for a year prior to this study, we excluded the 13 respondents who said they had seen and remembered this video (question 13) from all analyses.

The first outcome of interest in this research is whether the Star Trek-themed video was an effective way to teach people about prions. We tested this hypothesis in two ways: directly through question 5 above as well as indirectly through the difference in response between question 3 and question 2 above. This combination provided a secondary confirmation and allowed us to test our outcome of interest in two ways using both the stand-alone item and a calculated score. Research has demonstrated that asking people to retrospectively evaluate how much they knew about a topic before and after a learning experience is an accurate way to measure learning gains (Lang and Savageau, 2017). For the indirect measure, we calculated learning gains by subtracting the prior knowledge score from the subsequent knowledge score. Finally, we removed responses in which people said they already knew a lot (score of 4: 9 respondents) or a great deal (score of 5: 11 respondents) since those participants would be unable to experience meaningful learning gains by watching an introductory-level video.

Our second outcome of interest is whether the Star Trek connection helped make the videos more interesting and engaging for the participants. This outcome was central to our question of whether people would engage with science through popular culture connections. We measured this by directly through question 4.

Our third outcome of interest, related to the second, is whether the science fiction connection is broadening the viewership of this video content. Essentially, we sought whether some viewers would have been unlikely to watch a video on prions that did not relate to science fiction, and this was addressed with question 6.

Finally, we were interested in exploring how much people’s interest and engagement with the science video was affected by various parameters. The first was how familiar they were with the specific popular culture tie-in referenced in the video. In this study, we measured that possibility by asking people to indicate whether they were familiar with the *Star Trek: Discovery* episode that the video discussed (question 12). Only 7 respondents selected not knowing anything at all about Star Trek, so these responses were omitted from the test due to low sample size. We used this measure to test whether people’s engagement with the video varied based on their prior knowledge of the popular culture referenced in the video.

We also explored whether the venue and timing of most recent science education affected people’s engagement (question 10). Although 8 categories were presented, we excluded responses in 4 categories because of low sample size: never having had a formal science class (N=1), elementary or middle school (N=0), in the military (N=2), and through an online platform (N=12). This exclusion left the categories of high school (N=33), 2-year college or training program (N=24), 4-year college (N=98), and university graduate program (N=51). We also explored whether how long ago that class was affected engagement (question 11).

### Statistical analyses

Given likely non-normality of the data, non-parametric tests were used. To assess whether engagement in the content depended upon degree of familiarity with the series or episode, a Kruskall-Wallis test was conducted comparing responses regarding engagement in the video for respondents who are familiar with the Star Trek franchise vs. those familiar with *Star Trek: Discovery* but not the specific episode vs. those familiar with the *Star Trek: Discovery* episode. We used a Mann-Whitney U-test to then further explore whether those who remembered the episode were more engaged if they specifically remembered the role of prions in it. We used a Kruskall-Wallis test to explore if respondents with different levels of education exhibited different levels of engagement in the video. Finally, we used linear regression to assess if how long ago the most recent science class taken was associated with engagement.

Many more assessments or analyses were possible, but the few specific analyses and approaches described above were selected *a priori* so as to avoid statistical issues associated with multiple comparisons as well as “p-hacking”.

All survey questions, raw data, and limited dataset following exclusions will be archived in Dryad (datadryad.org) upon acceptance of this manuscript.

## RESULTS

### Self-reported learning

Respondents reported that the video increased their knowledge about prions. Figure 1 presents the counts and percentages, showing that over 88% of respondents reported at least “some” and over 99% reported at least “a little bit” of knowledge increase.

**Figure 1:**
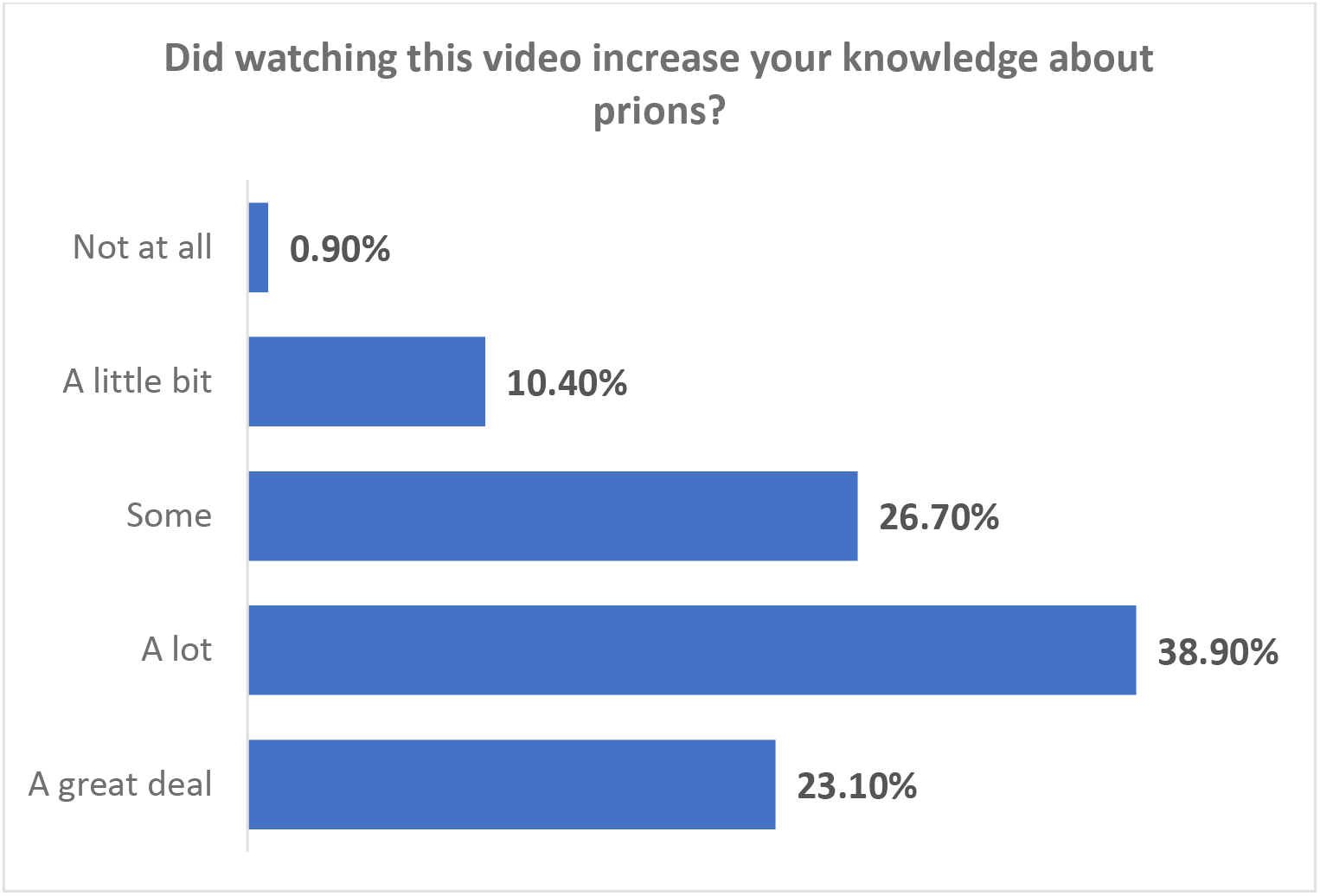

Similarly, when we subtracted the responses for how much respondents identified knowing about prions after watching the video minus before watching the video, only 7% (15) indicated no learning, and none reported knowing less. All the rest reported higher knowledge after watching the video relative to before watching the video.

### Science fiction connection and engagement/ interest

Respondents also overwhelmingly reported that the connection to the Star Trek franchise increased their engagement (figure 2). Only 10% reported no effect or that the Star Trek connection reduced their engagement.

**Figure 2:**
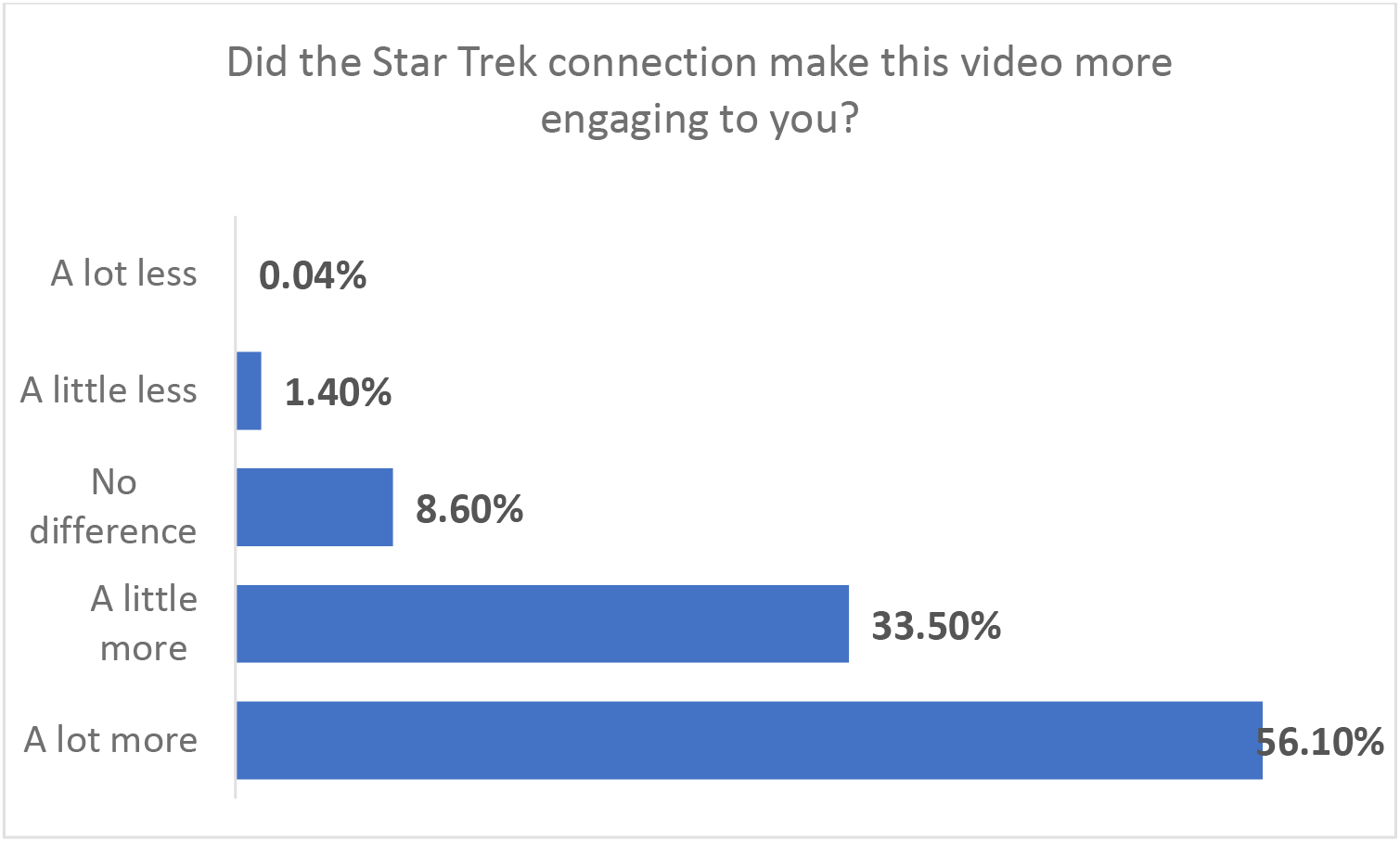

### Science fiction connection broadening viewership

Relatedly, several (48%) reported that they were “somewhat” or “very” unlikely to engage with such content if there were no Star Trek connection (figure 3). This fraction would likely be a minimum estimate of the subset of these viewers who were drawn to this science content primarily for the connection to Star Trek.

**Figure 3:**
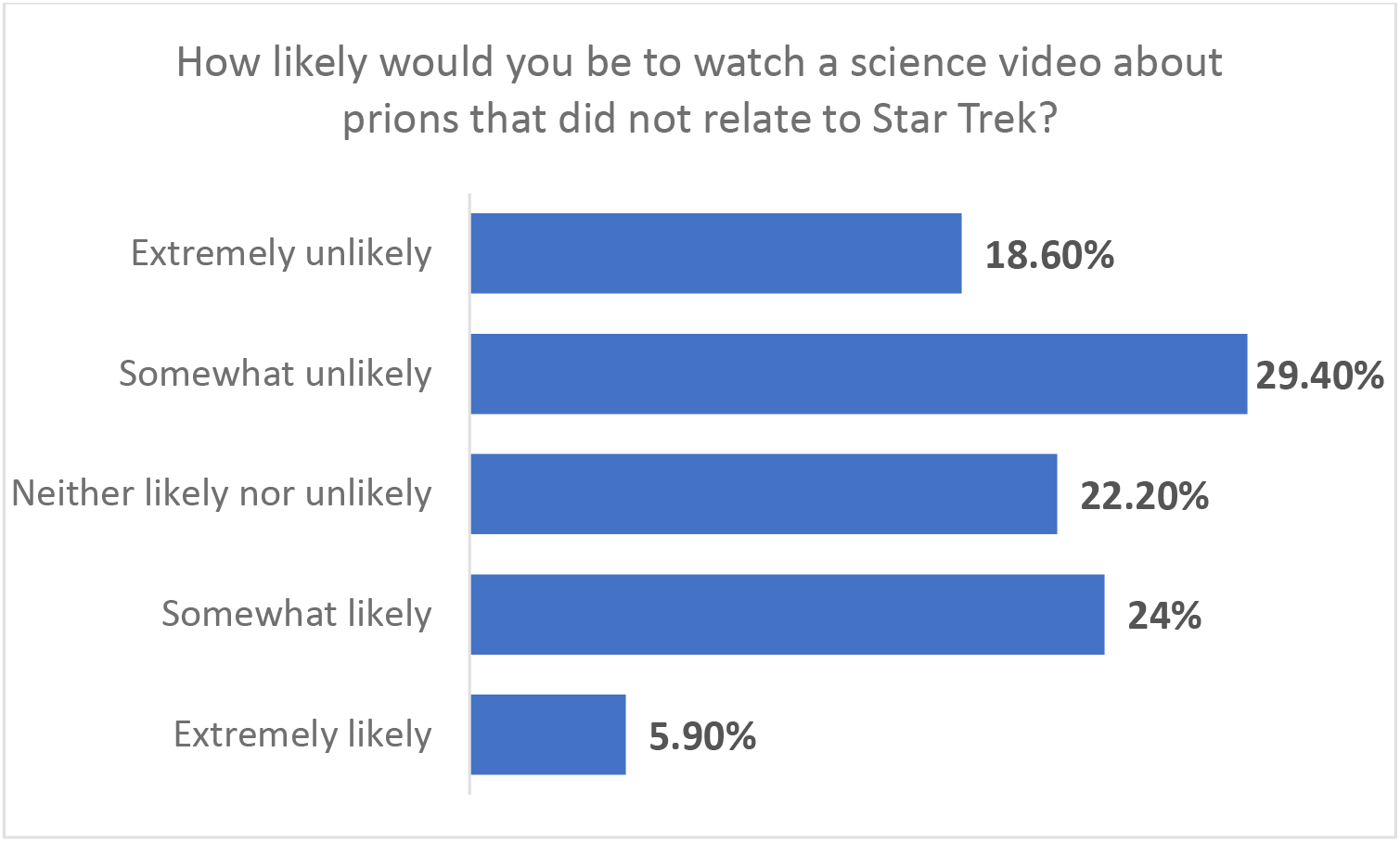

### Dependencies on science fiction franchise knowledge and education

We explored what parameters affected the self-reported engagement in this material: knowledge of the series or episode in question, level of education, or time since last formal science education. Knowledge of the series did seem to be associated with level of engagement, with individuals who know the series *Star Trek: Discovery* exhibiting on average higher levels of engagement than those who only knew the broader franchise (Kruskall Wallis test, 3 df, H=6.54, p=0.038). Only 4 people in total reported any negative effects on engagement because of the Star Trek connection, and these were spread among the categories of knowledge of the series. In contrast, 40% of the respondents who know Trek but not Discovery, 58% of those who know Discovery but not the episode, and 61% of those who know Discovery and the episode reported that the Trek connection made the video “a lot more engaging”. Specifically recalling the prion connection in the episode had no detectable effect (Mann-Whitney U-test, Z=0.69, p=0.49). However, we failed to detect any association between either venue of formal science education (Kruskall-Wallis test, 3 df, H=1.82, p=0.61) or time since last for formal science education (r=-0.002, p>0.5) affected engagement with the video.

## DISCUSSION

Our analysis found that the fictional narrative structure of an episode of *Star Trek: Discovery* created a pathway for people to engage with a science subject that they otherwise may not have sought out. This result demonstrates that using fictional stories from popular culture to disseminate science knowledge is both effective and engaging, particularly when the learners have a pre-existing interest or attachment to the popular culture works being leveraged. Further, we found that people need not have a strong attachment to the specific work of fiction being referenced. In our study, even people with only a general knowledge of the Star Trek franchise reported that the video was an attractive and effective way to learn about prions.

Taken together, our findings further support the use of fictional narrative connections to engage with non-student learners around science topics, extending the findings from classroom settings (Marsh et al., 2012) to a public online venue. Connecting science education with fictional narratives can make science content more attractive for at least a subset of learners and potentially may help them retain more of the content through the connection with popular culture, though we did not test this latter hypothesis.

Importantly, there is no *a priori* reason to assume that the results here are specific to the study topic (prions, or even biology) nor the science fiction franchise. Given the challenges associated with public science education, this approach of leveraging interest in science fiction to broaden science education outreach via educational videos represents a promising additional pathway for engagement.

Some caveats merit mention here. First, this qualitative study uses subjective self-reports on knowledge gains and interest. If we assume respondents are truthful in having watched the video per the instructions, at least some knowledge gain seems likely, but quantifying such a gain was not the focus of this study. Instead, the findings that 1) at least a subset of viewers found this mode of learning engaging and 2) a subset of viewers would have been unlikely to engage with similar science content without the science fiction connection are more relevant and novel. Second, this study does not attempt or purport to study a representative sample of the broader public. Based on how respondents were obtained, many of the subjects likely were positively predisposed to this particular popular culture franchise. One must not overinterpret the impact of the approach studied here: the same video may be much less effective at engaging the interest of a truly random sample of the public. Nonetheless, the goal of the study was to assess whether existing interest and predisposition in a particular popular culture franchise could be leveraged by even a subset of viewers predisposed to that franchise to teach scientific content, and this goal was successfully accomplished. One can hypothesize based on these results that other popular culture franchises can be leveraged in a similar manner to engage a different subset of viewers in science content.

## ACKNOWLEDGMENTS

We thank Sarah Marion and Elizabeth Marsh for helpful comments on this manuscript. This study was supported by NSF grant DEB-2019789 to MAFN.

